# Metaviz: interactive statistical and visual analysis of metagenomic data

**DOI:** 10.1101/105205

**Authors:** Justin Wagner, Florin Chelaru, Jayaram Kancherla, Joseph N. Paulson, Victor Felix, Anup Mahurkar, Héctor Corrada Bravo

## Abstract

Along with the survey techniques of 16S rRNA amplicon and whole-metagenome shotgun sequencing, an array of tools exists for clustering, taxonomic annotation, normalization, and statistical analysis of microbiome sequencing results. Integrative and interactive visualization that enables researchers to perform exploratory analysis in this feature rich hierarchical data is an area of need. In this work, we present Metaviz, a web browser-based tool for interactive exploratory metagenomic data analysis. Metaviz can visualize abundance data served from an R session or a Python web service that queries a graph database. As metagenomic sequencing features have a hierarchy, we designed a novel navigation mechanism to explore this feature space. We visualize abundance counts with heatmaps and stacked bar plots that are dynamically updated as a user selects taxonomic features to inspect. Metaviz also supports common data exploration techniques, including PCA scatter plots to interpret variability in the dataset and alpha diversity boxplots for examining ecological community composition. The Metaviz application and documentation is hosted at http://www.metaviz.org.

## Introduction

High-throughput sequencing of microbial communities provides a tool to explore associations between host microbiome and health status, detect pathogens, and identify the interplay of an organism’s microbiome with the built environment. Recent highlights include work on personalization of the human skin microbiome (Oh et al., 2014), diversity in the ocean microbiome (Sunagawa et al., 2015), and cataloging the global virome (Paez-Espino et al., 2016). As data is continually generated, developing analysis techniques and appropriate statistical models remains vital to gain insight from these projects. In other high-throughput sequencing assays, including human whole genome and transcriptome sequencing, software systems such as genome browsers that integrate exploratory computational and visual analysis have proven to be effective in analyzing these datasets (Chelaru, Smith, Goldstein, & Bravo, 2014; Kent et al., 2002). Extending this work to metagenomics presents challenges as metagenomic features, the units of measurement and analysis, do not map to the linear structures of tracks and ranges used in genome or gene expression visualization. As the features in metagenomic datasets have a hierarchy derived from annotation databases that link sequences to taxonomic classifications of bacteria, we use this hierarchy to build a navigation tool for effective exploration, analysis, and data visualization.

### Motivation

As an illustrative use case for statistically guided interactive visualization, we consider analyzing data from the Moderate to Severe Diarrheal disease study among children in four countries. Specific details for data generation, preprocessing, and annotation are covered in Pop *et al.* (2014). Raw count data is publicly available in the *msd16s* Bioconductor package [http://bioconductor.org/packages/msd16s/].

A typical analysis for this case-control study includes testing to compare taxa abundance between children with and without diarrhea to find novel associations with health and disease. The *metagenomeSeq* Bioconductor package [http://bioconductor.org/packages/metagenomeSeq] is a popular tool to identify differentially abundant features (Paulson *et al.* 2013). The first step of this approach is aggregating counts to a level of a taxonomic hierarchy (e.g. species or genus) and computing log fold-changes and p-values for each taxa between case and control groups. Then a selection can be made to retain only features with a log fold-change beyond a given threshold and p-value cutoff. The abundance of these filtered features across samples can then be visualized in a heatmap. After interpreting the plot, changing the feature selection or exploring the hierarchy requires another iteration of computing the feature set and generating a heatmap. Each refinement of parameters produces another visualization with no linking between results.

Our design of the Metaviz application for interactive visualization and analysis makes this workflow much more effective: once the set of features is selected, those changes can be propagated onto a web-browser visualization workspace. A user can then explore the hierarchy of features, aggregate counts to any level of the taxonomy, and identify sub-structures that are difficult to ascertain at lower levels of the taxonomic hierarchy. Further, differential abundance can be calculated at another level of the hierarchy then dynamically pushed to the same Metaviz workspace, thus streamlining the exploration of a complex set of differential abundance results between statistical and visualization tools.

## Design

We present the architecture of Metaviz from the web-browser application to database storage. A web-browser based application provides flexibility for users and “run anywhere” functionality when deploying the tool. We built upon the D3.js project for an aesthetically pleasing and effective suite of plots and charts. The back end serves an abundance matrix with taxonomic annotation for features, in our case Operational Taxonomic Units (OTUs), and the front end is a JavaScript application for data visualization. Given the structure of metagenomic data, the user navigation tools and the database storage are tailored to taxonomic hierarchies. We moved from a relational database model used in Epiviz (Chelaru & Bravo, 2015), an interactive application for visualization and analysis of functional genomic data such as gene expression and methylation data, to a graph database to hold the feature hierarchy and abundance counts. A fundamental operation we enable is specifying a set of nodes in the hierarchy of features and aggregating counts to that set.

### Visualization layer

Implementing the visualization layer presents several challenges for displaying, navigating, and manipulating data from a feature-rich hierarchy. Design considerations for metagenomic data analysis include: 1) *size of the feature space,* which in datasets we visualized using Metaviz, ranges from 47 (unpublished collaborator data) to 45,000 (Human Microbiome Project) features; 2) *depth of the feature hierarchy,* which is a function of the annotation database; and 3) *number of samples,* with as many 992 (*msd16s*) samples in a dataset we analyzed. Given these characteristics, we focused the design of Metaviz on efficient traversal of the feature space and defining feature selections across the taxonomy. In addition, we engineered the navigation tools to be applicable across datasets and persistent between user sessions for collaboration and publication of results.

In Figure 1, we demonstrate the visualization layer of Metaviz on a metagenomic dataset. The bottom panel is a novel navigation control designed to effectively explore the taxonomic feature hierarchy and aggregating count values of features to any set of taxonomic nodes. The top panel consists of a heatmap with the color intensity set as the observed count of a feature (column) in a sample (rows). The rows are dynamically clustered based on Euclidean distance of the count vectors for each sample and a dendrogram shows the clustering result. The top panel also includes a PCA plot over all the features of the samples in the heatmap. The stacked bar plots in the second row render, for each sample (column), the proportion of counts for each bacterial feature. The separate plots show case (left) or control (right) samples based on dysentery status and the columns are samples grouped by age range. The collection of charts provides multiple views of the same data and are dynamically updated upon user interaction with the navigation tool to achieve exploratory iterative visualization.

**Figure 1:**
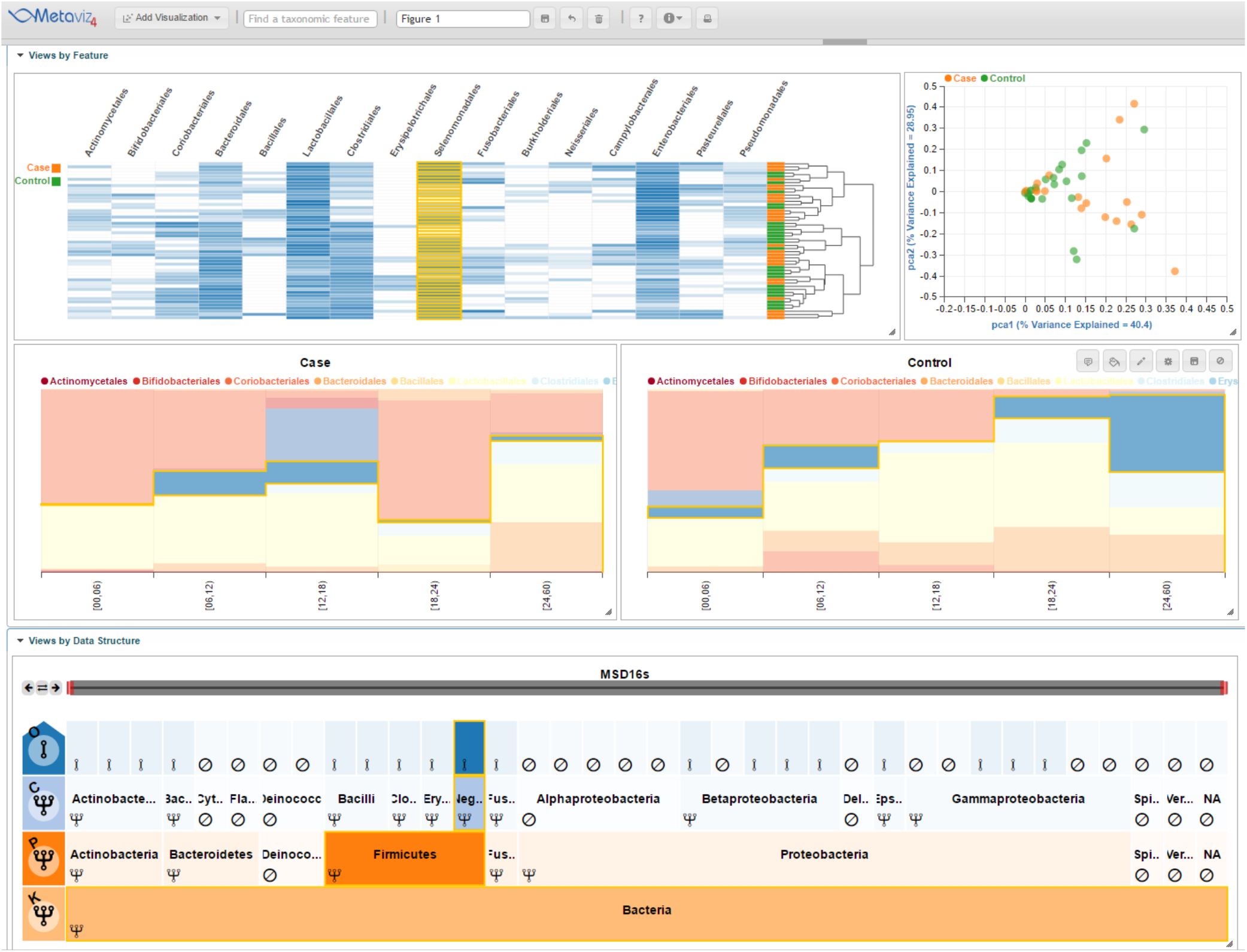
Metaviz interactive visualization of childhood severe diarrhea study. This figure shows a subset of 50 samples (25 case and 25 control for dysentery) from the Moderate to Severe Childhood Diarrheal Disease study (Pop et al., 2014). The FacetZoom control on the bottom panel is used for interactive exploration of the taxonomic organization of metagenomic features. Node opacity in the FacetZoom control indicates the set of taxonomic units selected across all appropriate visualizations in the Metaviz workspace. Each node can be in one of three possible states as indicated by the icon in its lower left corner: 1) *aggregated,* where counts of descendants of this node are aggregated and displayed in other charts, 2), *expanded,* where counts for all descendants of this node are visualized in other charts, or 3) *removed,* where this node and all its descendants are removed from all the other charts. The left column of the FacetZoom control indicates the levels of the taxonomy and the overall selection for nodes at each taxonomic level. Hovering the mouse over FacetZoom panels highlights the corresponding features in other charts through brushing. The top left chart is a heatmap showing log-transformed counts with color intensity in each cell corresponding to the abundance of that feature (column) in that sample (row). The row dynamically computed and rendered dendrogram shows Euclidean distance hierarchical clustering of samples with color indicating case/control status of each sample. The yellow highlighted column is linked between charts and FacetZoom control through brushing. The top right chart is a PCA plot over all features for the samples selected. The stacked bar plot on the left of the second row shows proportion of selected features in each case sample (columns) while the right chart shows control samples. In both, sample counts are grouped and aggregated by age range. This is available as a Metaviz workspace at http://metaviz.cbcb.umd.edu/?ws=yA4BWgUOTiq.

#### Using Information Visualization Techniques for Metagenomics

Our design for the visualization layer is motivated by results in the information visualization literature for displaying large tree structures with associated complex data. In this section, we provide a brief review of pertinent visualization techniques. To provide a basis for our design decisions, we present metagenomic visual analysis operations in relation to the Task by Data Type Taxonomy for Data Visualization (Shneiderman, 1996). In metagenomic sequencing projects, sample data is multi-dimensional with study-specific attributes, e.g. age, sex, gathered in each experiment. Feature data is tree-structured with a node fan-out dependent on the bacterial hierarchy of the annotation database and the ecological community observable in each sample.

We review the tasks presented by Shneiderman for completeness. These consist of the following: 1) *Overview:* gain an overview of the entire collection; 2) *Zoom*: Zoom in on items of interest; 3) *Filter*: filter out uninteresting items; 4) *Details-on-demand*: Select an item or group and get details when needed; 5) *Relate*: View relationships among items; 6) *History*: Keep a history of actions to support undo, replay, and progressive refinement; 7) *Extract:* Allow extraction of sub-collections and of the query parameters (Shneiderman, 1996). Our task taxonomy below builds upon and generalizes the description of features presented in the Krona interactive visualization tool (Ondov, Bergman, & Phillippy, 2011), also based on the Shneiderman interactive visualization task taxonomy.

We now discuss the specific operation and goal for each task with regards to metagenomic analysis. The *overview* task consists of examining global patterns in feature abundance among samples across levels of the taxonomic hierarchy. This task is also accomplished by presenting statistics that summarize feature variance and observed ecological diversity. The *zoom* task requires navigation to the lowest levels of the feature hierarchy as well as inspection of individual sample data. The *filter* task consists of removing or expanding taxonomic features and samples. With metagenomic data, several operations need to be enabled, first a level-wise filtering and then removal of features at a given depth along with aggregating to a specific point in the hierarchy. *Details-on-demand* includes showing all children of a given node, text-based search for features that contain a character string, and the utility to visualize the same data in different views. *Relate* in metagenomics is enabled by linking multiple data views with the feature hierarchy along with group-by and color-by operations over sample attributes. *History* requires keeping track of the current position during navigation of a feature hierarchy as well as the ability to select and remove nodes as desired. Finally, *extract* entails capturing the parameters to recreate an analysis. Specific to metagenomic analysis, the *extract* task also should encompass providing a mechanism to interoperate between annotation databases and retrieving cluster center sequences from a dataset.

#### Navigation Mechanism - FacetZoom

We developed a novel navigation technique to handle the complex and hierarchical structure of metagenomic feature data that enables the visualization tasks of *overview, zoom,* and *filter.* The design is inspired by aspects of an icicle plot, which shows all children under a given node and was historically proposed for biological taxonomy visualization (J. B. Kruskal, 1983). We also incorporate a technique, FacetZoom, that visualizes a hierarchy using a tree structure showing a subset of levels at one time (Dachselt, Frisch, & Weiland, 2008). Our navigation tool is designed to handle the limitations in the screen size and performance of rendering trees with tens of thousands of nodes. We refer to our navigation utility, shown in the bottom panel of Figure 1, as a FacetZoom control for the rest of the manuscript.

The nodes of the FacetZoom control indicate how the abundance counts for taxonomic features are displayed in the charts of the Metaviz workspace. Every node of the FacetZoom control can receive mouse-click input from the user. A click on a node sets that feature as the root of a dynamically rendered subtree. Each node can be in one of three possible states as indicated by an icon in its lower left corner: 1) *aggregated,* where counts of descendants of this node are aggregated and displayed in other charts, 2) *expanded,* where counts for all descendants of this node are visualized in other charts, or 3) *removed,* where this node and all its descendants are removed from the other charts. The state of a node is propagated to all its descendants. Node opacity in the FacetZoom control indicates the set of taxonomic units selected across all appropriate visualizations in the Metaviz workspace. Hovering the mouse over FacetZoom nodes highlights the corresponding features in other charts through brushing as shown in Figure 1.

The FacetZoom control includes a level-wise aggregation indicator panel on left side. Each element of the indicator panel provides information on the current depth in the hierarchy and can be used to set the state of all nodes at a given depth. The letter on the indicator panel identifies the taxonomic level with “P” denoting phylum and “O” signifying order, for instance. The top and bottom nodes of the indicator panel display the directions in the hierarchy containing additional levels to visualize.

The bar at top of the FacetZoom sets the range of features to query and visualize. The bar is a flexible component with arrows to control movement left or right and expansion over the full range of the current hierarchy. Updates to the filter bar triggers queries over the count data and those results are automatically propagated to the other charts in the workspace.

#### Data Plots and Charts

We provide several visualizations of feature count data. These allow the user to explore relationships between sample phenotype and metagenomic features. The first is a heatmap with rows as samples and columns as features (Sneath, 1957). The heatmap is an interactive component from which a user can select to show a dendrogram of a dynamic clustering over features or samples. If the user chooses not to employ clustering, rows can be re-ordered based on a sample metadata attribute. We also provide several utilities on the samples including color-by and modifying the displayed name of any sample attribute. Figure 1 shows a heatmap of the *msd16s* dataset with the colors for sample rows set based on dysentery status.

Another visualization in Metaviz is the stacked bar plot that shows the proportion of features in a given sample. On the stacked plot, we implemented a group-by function to aggregate samples based on a sample metadata attribute. This plot is useful for comparing microbial community composition between individual samples or groups. Figure 1 shows two stacked bar plots that are split based on sample dysentery status and grouped by age range.

Metaviz supports scatter plots to visualize feature count values of selected samples in a X, Y coordinate plane. A scatter plot is useful for fast identification of distribution and spread across measurements. The scatter plot has a color-by feature to color points based on a specific sample metadata attribute. One specific scatter plot is the PCA plot seen in the upper left corner of Figure 1.

Further, Metaviz includes a line plot with each line representing a feature, the height of the line denoting abundance, and the samples across the X-axis. We find the line plot useful for examining time-series data. Metaviz also provides a box-plot for visualization of alpha diversity with each box as a group of samples with a selected attribute.

All data plots and charts added to the workspace are linked to the feature nodes on the FacetZoom. Hovering over a feature column in a heatmap highlights that feature in all other plots as well as the path through the hierarchy for that feature in the FacetZoom. This brushing and linking is essential to providing integrative visual analysis. Also, each plot and chart has a toolbar that can be used to modify presentation settings, the color scheme, saving to an SVG, and writing custom JavaScript for that chart. The toolbar is shown in the upper right hand corner of the stacked plot for control samples in Figure 1.

#### Text Search

Metaviz supports text-based search for quick navigation to specific taxonomic features. A user can enter a feature of interest into a search box on the toolbar. The search provides auto-complete and a list of features that contain the character string are displayed in a drop-down list. Once a user selects a feature, the filter bar in the FacetZoom control will update to encompass that feature and all linked data visualizations update as well.

### Data layer

A key difference between metagenomic data and other genomic data is the hierarchical organization of its features, which drives the design of the Metaviz back end. Our data model of metagenomic datasets includes the observed counts for each feature in every sample, the hierarchical taxonomic feature annotations and metadata such as phenotypic, behavioral, and environmental information for each sample. A query triggered from the UI operates over these three data types and computes aggregation on the count data to the specified hierarchy level.

#### Graph Database

Our initial backend architecture separated the counts, sample metadata, and feature hierarchy into relational database tables. An aggregation query operated on the feature hierarchy table followed by a join with the sample and count tables to arrive at the result. This approach worked for developing a proof-of-concept, but to achieve interactive visualizations with reasonable query response times, we ultimately chose to implement a graph database architecture.

In a graph database, nodes and edges in a graph are first class objects that can be queried directly. This is a contrast to relational databases in which samples are rows and sample attributes are columns. Each table in a relational database encompasses all the required data fields for the observations in that table while keys handle relationships between tables. As the join operation is the costliest operation in our three-table design, we use a graph database to store each feature as a node and each feature has an edge connecting it in the taxonomic hierarchy. We also store samples as nodes and the count value for a feature in a sample is an edge between leaf feature nodes and sample nodes. This graph database structure is shown in Figure 2.

**Figure 2:**
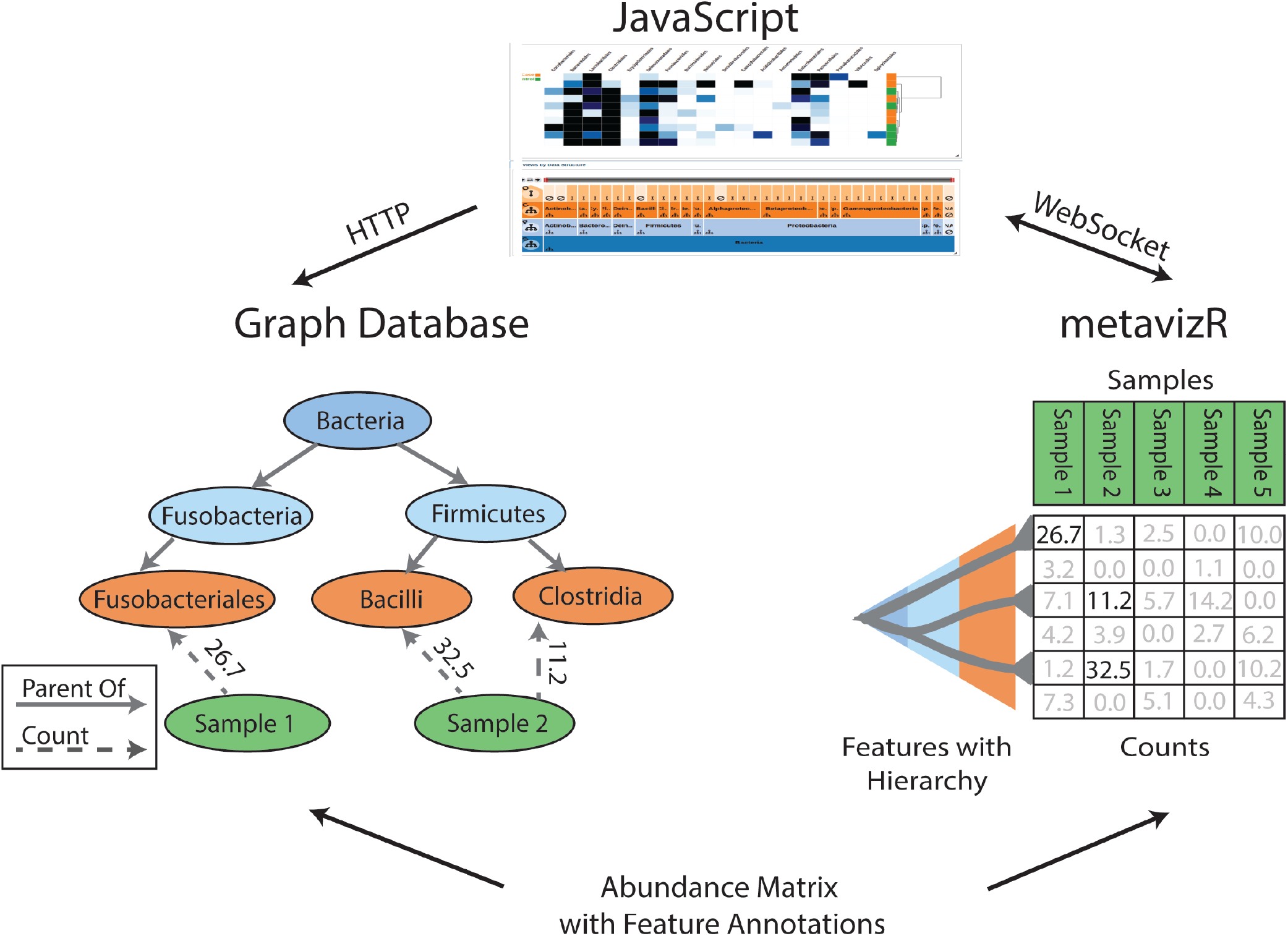
Metaviz query processing and Graph DB structure. There are two deployment options, which can be used concurrently if desired. In one deployment option (left), the Metaviz JavaScript front end makes requests to a Python application querying a graph database using HTTP. In the other deployment option (right), abundance matrices are loaded into a *metavizr* session which uses the WebSocket protocol to communicate to the JavaScript component, allowing two-way communication between JavaScript and an interactive R session. The graph on the left shows how abundance matrices are stored in the graph database. Nodes in the graph correspond to metagenomic features or samples, edges between metagenomic features denote taxonomic relationship, edges at the leaf level of the taxonomy connect to samples and store the corresponding abundance counts. In either deployment option, aggregation queries are evaluated in response to FacetZoom control selections in the UI and require summing, for each sample, the counts for features in a selected taxonomic subtree.

To inform the choice of graph database versus relational database, we benchmarked an implementation of the three table design using MySQL-PHP against a Neo4j-Flask deployment. In the benchmarks, we deploy our backend services on an Amazon EC2 t2.small instance and used the *wrk* tool [https://github.com/wg/wrk] to send HTTP requests. The testing dataset consisted of 62 samples, 973 features, and 7 hierarchy levels. We observed that compared to our initial MySQL-PHP approach, the graph database provides 5x lower latency. We also modified our MySQL-PHP design to pre-compute the join operation across the three tables and store that in the database as well. Compared to this implementation, our Neo4j-Flask implementation showed approximately 50% lower latency. We show our benchmark results in Figure 3.

**Figure 3:**
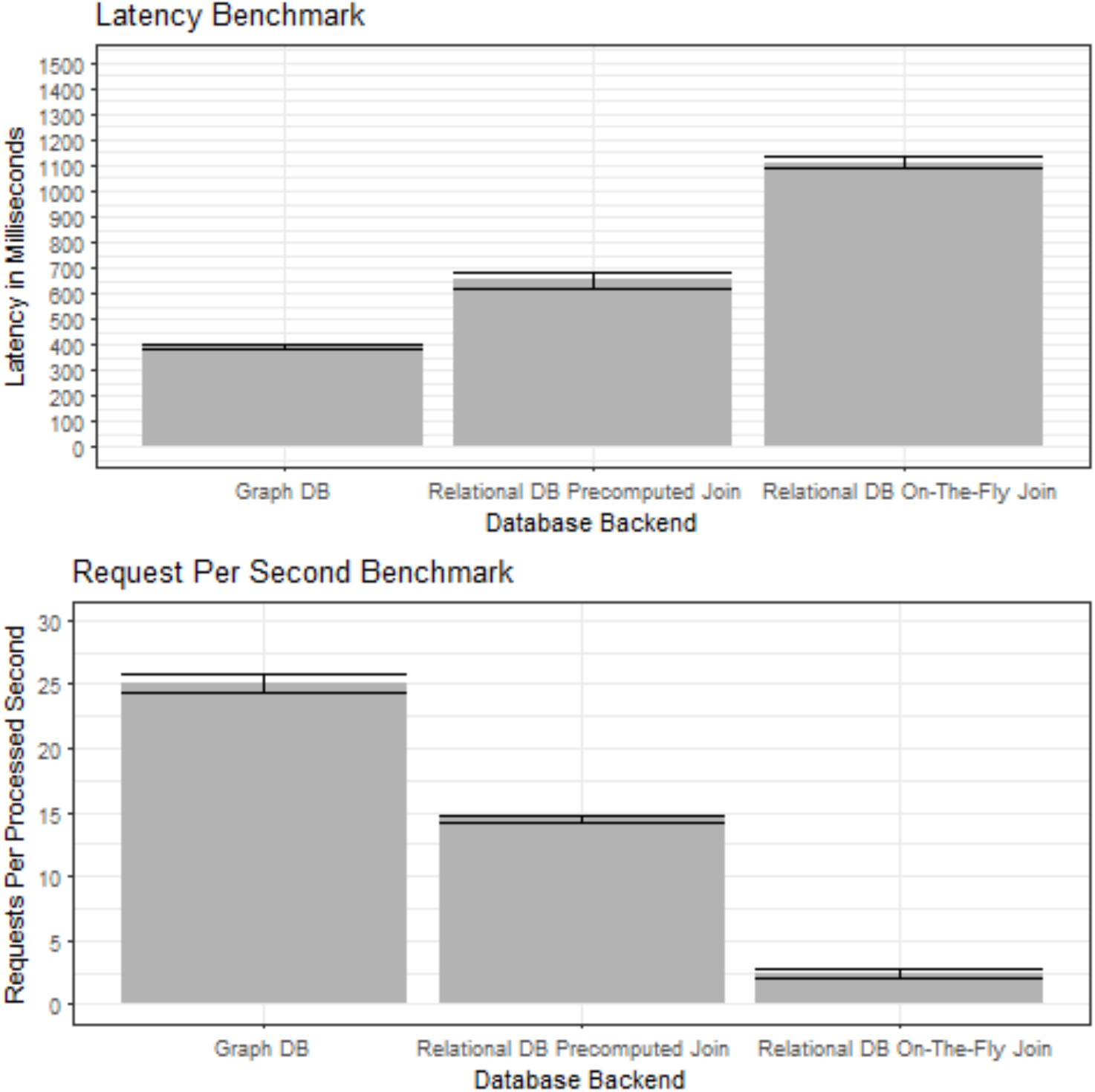
Metaviz database architecture benchmarks. We use the *wrk* [https://github.com/wg/wrk] tool to benchmark UI requests to three database architectures for storing abundance matrices and feature hierarchies (taxonomies): (1) Graph DB, using Neo4j with a Python Flask web service, (2) Relational DB Pre-computed Join, using a MySQL implementation with a JOIN of the 3 tables of features, values, and samples pre-computed and stored as a table, (3) Relational DB On-The-Fly Join, a MySQL implementation with computing a JOIN across the three tables for each query. For (2) and (3), a PHP application issues queries to the database in response to requests from the UI. We deployed each implementation on an Amazon EC2 t2.small instance and the dataset used across all instances consisted of 62 samples, 973 features, and 7 hierarchy levels. The upper panel shows query latency including standard error across 5 days of measurements. In addition to the latency of processing each request, we also measure the number of requests per second processed providing a measure of throughput in our application. In both performance measures, we see significant benefits of a Python-Neo4j deployment compared to a PHP-MySQL stack for Metaviz tasks.

#### metavizr

In addition to the persistent database backend, Metaviz can also perform interactive visualization with the *metavizr* R package [https://github.com/epiviz/metavizr]. *Metavizr* loads an *MRexperiment* object, the main class of the *metagenomeSeq* R/Bioconductor package for statistical analysis of metagenomic data, into an *EpivizMetagenomicData-class* object. This R object can then communicate with a Metaviz application instance using a WebSocket connection. FacetZoom controls along with data charts and plots can be added to the UI from the R session. A user can specify features for visualization from the results of statistical testing as discussed in the *Motivation* section. The feature selection can use R/Bioconductor packages beyond *metagenomeSeq.* For instance, we use the *vegan* CRAN package to compute alpha diversity in *metavizr* sessions (Oksanen et al., 2015). Github gists can be used through *metavizr* to modify any plot or chart display setting using JavaScript in addition to customization facilities provided directly by the *metavizr* package itself. Finally, a persistent workspace can be used to reproduce the visual analysis of another researcher after *metavizr* loads the dataset.

To measure the performance of *metavizr* relative to the graph database architecture, we benchmarked the memory usage and run-time of aggregation operations on datasets of different sizes. Figure 4 show the profiling results. From these benchmarks, we recommend using the graph database backend for abundance matrices that are larger than 4000 features and 100 samples.

**Figure 4:**
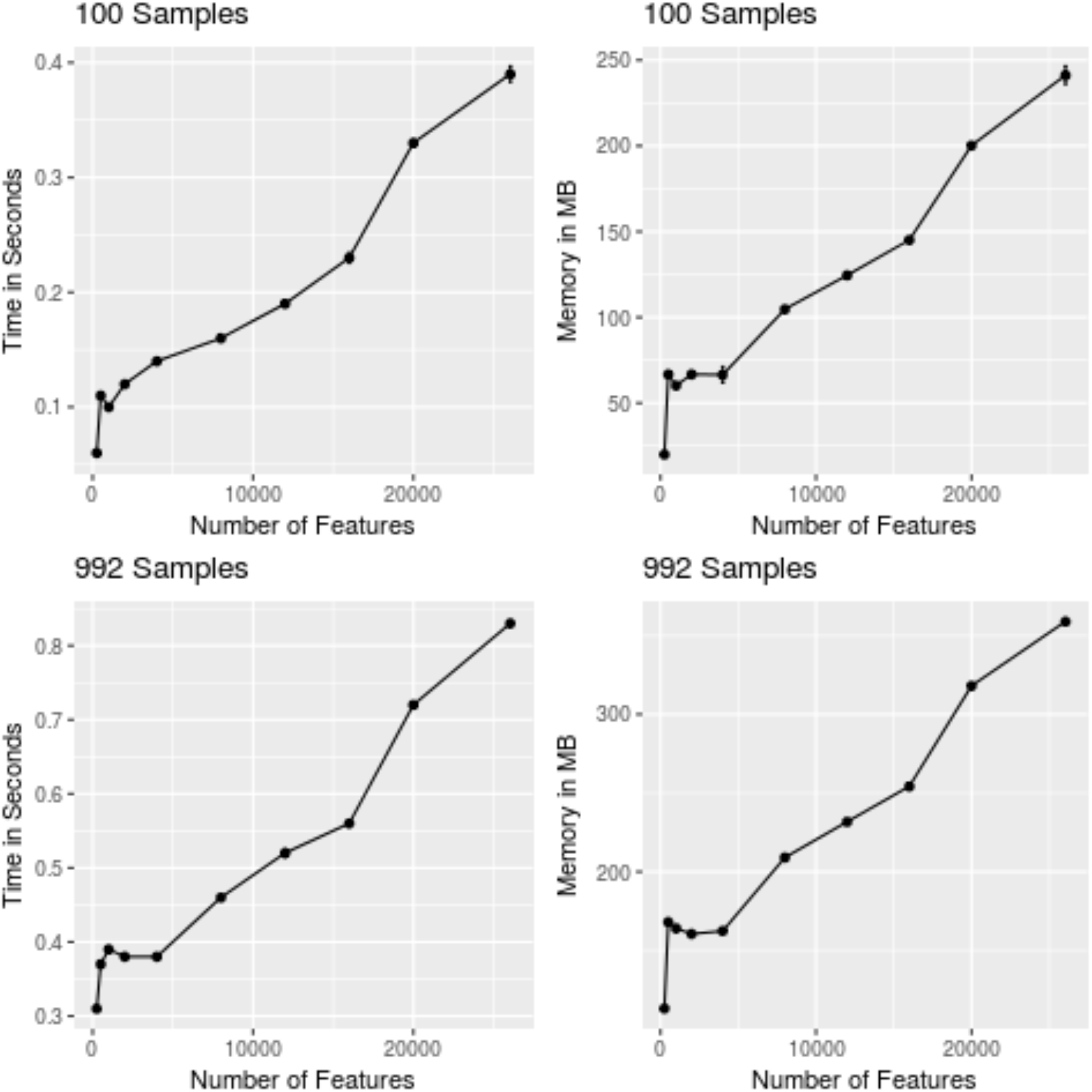
metavizr benchmark. MSD16s dataset with 992 samples, 26044 features, and a 7 level hierarchy. The Rprof library was used for profiling. The benchmark consisted of an aggregation query to the 3rd level of the hierarchy. The top panels show tests for keeping the number of samples at 100 and increasing the number of features over which the aggregation query is operating. The left top panel shows the aggregation query completion time in seconds and the top right panel shows the highest memory footprint in MBs during the query execution. The bottom panels show the performance on all samples in the dataset and increasing the number of features. The bottom left panel shows the relationship between aggregation query time and number of features while the bottom right panel is the memory footprint in MBs use during the aggregation. In order to achieve interactive visualization, we recommend using the Neo4j graph database when aggregation query response times will be above 400 milliseconds. Also, the graph database backend is recommended for datasets that require a memory footprint above 200 MB.

### Deployment

We support three deployment mechanisms of Metaviz for users to interactively visualize an abundance matrix with hierarchical feature annotations. The first scenario is hosting the Metaviz browser application and graph database locally. We provide Docker [http://www.docker.com] scripts to build and deploy containers of the database, load the abundance matrix to the database, and launch the web-browser application [https://github.com/epiviz/metaviz-docker]. The second deployment option is loading the abundance matrix into an *MRexperiment* object with the *metagenomeSeq* R/Bioconductor package and creating a *metavizr EpivizMetagenomicsData-class* object. Then, *metavizr* can communicate with the client’s web-browser through an R session. We also support hosting datasets at the University of Maryland Center for Bioinformatics and Computational Biology. The abundance matrix will be loaded into a Neo4j database and can be accessed from http://metaviz.cbcb.umd.edu. A user can perform analysis and then share results with collaborators using the persistent workspace functionality.

## Use cases

### Exploration of MSD16s childhood diarrhea study in developing countries

To test the analysis utility of Metaviz, we inspected the *msd16s* dataset using the web-browser application. To visualize and explore the samples, we examined the data from each country in the study separately and aggregated to the order level for all features. In this analysis, we set case samples as those with dysentery and control samples as those without blood in stool, meaning that samples with diarrhea and healthy samples are in the control group for dysentery. We used three plots, a heatmap and two stacked bar plots to identify differences between age range and dysentery status by country. In the heatmap, row colors were set by dysentery status and each stacked bar plot consisted of the case and controls samples for dysentery of each country. We also grouped the samples in the stacked bar plot by age range. We compared the results of visual analysis by computing the log fold-change using *metagenomeSeq* and identifying features detected as differentially abundant via the statistical method and those detected through visual exploration.

From the heatmap for Bangladesh samples, the following orders appear more abundant in the samples with dysentery than the control samples: Actinomycetales, Burkholderiales, Neisseriales, Campylobacterales, Pasteurellales, and Pseudomonadales. Correspondingly, these orders appear more abundant in the control samples as compared to the case samples: Coriobacteriales, Bacteroidales, and Clostridiales. Looking at the stacked bar plots, Clostridiales exhibits low proportion in the case samples at 0-6 and 6-12 months, a lower level compared to control samples at 12-18 months, and then a similar proportion in both groups for 18-24 and 24-60 months. With the control samples, Bacteroidales and unannotated (NA) show a greater proportion at all intervals after 0-6 months. Supplementary Figure 1 shows the visual analysis of Bangladesh samples.

We followed the visual analysis by differential abundance testing using *metagenomeSeq.* The following orders had a log fold-change above 1 and an adjusted p-value < .1 when comparing the dysentery samples against non-dysentery: Acidithiobacillales, Pasteurellales, Spirochaetales, Campylobacterales, Neisseriales, and Enterobacteriales. Through visual analysis we found Campylobacterales (log fold-change: 2.30, p-value: 2.46*10 ^-4), Neisseriales (1.92, 3.55*10 ^-5), and Pasteurellales (2.40, 4.96*10 ^-11). The orders identified from visual inspection but not through *metagenomeSeq* had the following log fold-change and adjusted p-values: Actinomycetales (log fold-change: .973, adjusted p-value: 7.65*10 ^-3), Burkholderiales (0.417, 0.400), and Pseudomonadales (0.834, 1.83*10 ^-2). These are the orders with a log fold-change below −1 and adjusted p-value < .1 in our analysis: Coriobacteriales (−1.27, 6.96*10 ^-4), Bacteroidales (−1.23, 3.77*10 ^-4), and Clostridiales (−1.20, 6.95*10 ^-5). We identified each of those orders through visual analysis. The results for *metagenomeSeq* differential abundance calculation are in Supplementary Table 1.

Building the same Metaviz plots for The Gambia, we note that the number of control samples outweighs the number of case samples and no case samples from the 0-6 month age range are present. Examining the heatmap, it appears that Actinomycetales, Lactobacillales, Campylobacterales, Enterobacteriales, and Pasteurellales are more abundant in the case than control samples. While Bacteroidales and Clostridiales are more abundant in the control samples than case. Examining the stacked plots, we first notice the proportion of Bacteroidales increases with age in the control samples as compared to the dysentery group. Lactobacillales decreases in proportion as age increases for both the case and control samples with a large decrease from 18-24 to 24-60 months in the case samples. In the case samples, Enterobacteriales has one of the highest proportions orders at 0-6 months, decreases for both 12-18 months and 18-24 months, but is then the highest proportion order in the 24-60 month interval. In the control samples, the proportion of NA increases at each age range. For case samples, NA shows lower proportion in 6-12 and 24-60 months than the 12-18 and 18-24 months intervals. Supplementary Figure 2 shows the Metaviz workspace for The Gambia.

Using *metagenomeSeq* we found the following orders to have a log fold-change above 1 and an adjusted p-value < .1 in case samples compared to control: Pasteurellales (1.54, 4.70*10 ^-4), Enterobacteriales (1.46, 1.26*10^-2), and Actinomycetales (1.13, 1.26*10^-2). We found Actinomycetales, Pasteurellales and Enterobacteriales through visual analysis. We also identified Campylobacterales (1.10, 0.133) and Lactobacillales (0.624, 0.377) while inspecting the heatmap but these were not differentially abundant using *metagenomeSeq.* Our visual analysis showed Bacteroidales (−1.76, 3.02*10^-3) and Clostridiales (−.833, .175) more abundant in control samples but *metagenomeSeq* analysis yielded only Bacteroidales. We present the *metagenomeSeq* differential abundance calculations for The Gambia in Supplementary Table 2.

Inspecting the Kenya samples with Metaviz we noticed there are far fewer samples with dysentery than non-dysentery samples. From the heatmap, we observed Actinomycetales, Lactobacillales, Fusobacteriales, Enterobacteriales, and Pasteurellales to be more abundant in the case samples than across the control samples. Bifidobacteriales, Bacteroidales, Coriobacteriales, and Clostridiales appear more abundant in control over case. As for changes across age ranges and case/control status, Campylobacterales is more prevalent in 0-6, 6-12, and 12-18 month age ranges in the case group than the control group. Supplementary Figure 3 shows the visual analysis of the Kenya samples.

Testing with *metagenomeSeq* showed Pasteurellales with a log fold-change above 1 and an adjusted p-value < .1 when comparing case against control samples. The Pasteurellales (<u>1.09, 3.76*10</u>^<u>-2)</u> result matched our findings from visual analysis but the Lactobacillales (.201, .949) Actinomycetales <u>(0.365, 0.572),</u> Fusobacteriales <u>(4.19*10</u>^<u>-2, 0.949), and</u> Enterobacteriales <u>(0.509, 0.430)</u> results did not. Using *metagenomeSeq,* we did not find any orders with log fold-change below −1 and adjusted p-value < .1. In Metaviz, Coriobacteriales (−.993, 7.31*10^-2), <u>Bifidobacteriales</u> (−3.50*10^-2, 0.949), Bacteroidales (−1.16, 0.146), and Clostridiales (−.898, .153) appeared to be more abundant in control samples than case samples. We show the *metagenomeSeq* differential abundance calculations for Kenya samples in Supplementary Table 3.

From the Metaviz plots for Mali samples, we note that the number of case samples is far smaller than the number of control samples and no case samples are from the 0-6 month age range. Examining the heatmap, Neisseriales, Lactobacillales, Enterobacteriales, and Pasteurellales show greater abundance in the case samples. In contrast, Bifidobacteriales, Bacteroidales, Clostridiales, and Pseudomonadales exhibit higher abundance in control samples compared to the case samples. From the stacked plots, the proportion of Enterobacteriales among case samples in age range 6-12 and 12-18 months is much higher than that in the same age ranges for control samples. For dysentery samples, Pasteurellales shows a much higher proportion in the 18-24 month age range than for normal samples. Also, across all age ranges Bacteroidales is more prevalent in the control samples. Supplementary Figure 4 shows the visual analysis of samples from Mali.

With *metagenomeSeq,* we observe Pasteurellales <u>(2.82, 5.39*10</u>^<u>-4)</u> and Neisseriales <u>(1.77, 1.38*10</u>^<u>-2)</u> to have a log fold-change above 1 and adjusted p-value < .1 for abundance in case against control. We found Pasteurellales and <u>Neisseriales</u> through visual inspection along with Enterobacteriales <u>(1.46, 0.799) and Lactobacillales (6.82*10</u>^<u>-2, 0.998).</u> From Metaviz visual analysis, we found Bifidobacteriales (−1.59, .439), Pseudomonadales (0.662, 0.460), <u>Bacteroidales</u> (−1.16, 0.439), and <u>Clostridiales (−1.80*10</u>^<u>-3, 0.998)</u> to show higher abundance in control samples but that was not borne out in statistical testing. Supplementary Table 4 contains the *metagenomeSeq* results for Mali samples.

### Analysis of longitudinal metagenomic studies

Another use case we sought to test the effectiveness of Metaviz is the analysis of longitudinal metagenomic datasets. We followed the analysis using smoothing spline ANOVA as described in Paulson et al. (Paulson, Talukder, & Corrada Bravo, 2017) for a longitudinal dataset understanding host response to a challenge with enterotoxigenic *E. coli* (Pop et al., 2016). The dataset was gathered from 12 participants who were challenged with E. *coli* and subsequently treated with antibiotics. Stool samples were gathered from participants each day starting 1 day pre-infection until 9 days post-infection. This time series analysis using smoothing spline ANOVA yields features that are differentially abundant across timepoints. The *metagenomeSeq* Bioconductor package provides a function for fitting the smoothing spline and performing SS-ANOVA testing. For visualizing the results of that analysis, we use a line plot with time points on the X-axis, log fold-change on the Y-axis, and each line representing a feature. The FacetZoom is linked to the line plot and the path through the hierarchy is highlighted when hovering over a given line. Figure 5 shows the visualization of the smoothing spline using *metavizr.*

**Figure 5:**
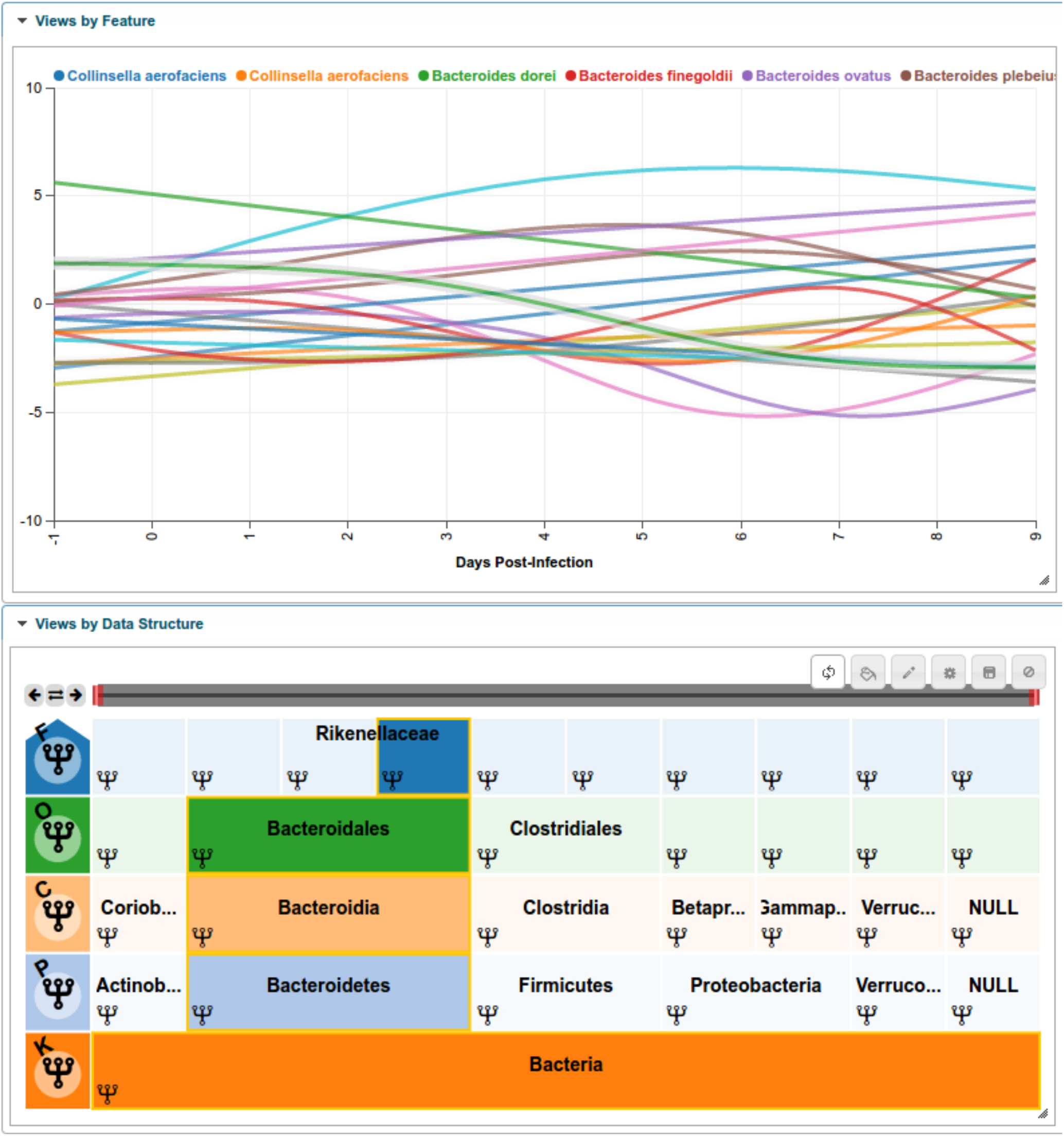
Interactive visualization of smoothing spline differential analysis of longitudinal study. We use Metaviz to explore a longitudinal analysis of the dataset from an enterotoxigenic *E. coli* study (Pop et al., 2016). Count data was aggregated to the species level and a smoothing-spline ANOVA model was fit using the *fit TimeSeries* function of the *metagenomeSeq* Bioconductor package. Features with a statistically significant difference of an absolute log fold-change greater than 2.5 as estimated by the smoothing spline model at any time point were selected for visualization. The line plot is linked with the FacetZoom control via brushing.

## Conclusion

In this paper, we presented the design and performance of a web-browser based interactive visualization and statistical analysis tool for metagenomic data. We described design decisions for operating over abundance matrices with tens of thousands of features, thousands of samples, and complex feature hierarchies. We use a graph database for storing metagenomic abundance matrices as the features have a hierarchy derived from taxonomic databases. We also built an R package for serving data to the Metaviz web-browser application and computing analyses using R/Bioconductor packages. A major contribution of this work is the novel navigation utility that adapts information visualization techniques to effectively explore and manipulate the rich feature hierarchy of metagenomic datasets. Another significant contribution is a web service available to host abundance matrices that allows researchers to explore and share results. We expect that Metaviz will prove useful for researchers in analyzing metagenomic sequencing studies as genome browsers have for genomic data.

### Related Work

We designed and implemented Metaviz while continually examining the list of successful projects for interactive visualization of sequencing data. The UCSC genome browser provides a track-based web tool for exploring the linear structure of genome and transcriptome sequencing data (Kent et al., 2002). Galaxy provides an integrated analysis environment and visualization tools that can be run in a hosted computing setup (Afgan et al., 2016). Taxonomer performs both read taxonomic assignment and visualization of results using a sunburst diagram to visualize features (Flygare et al., 2016). Pathostat is a Shiny application that computes statistical metagenomic analyses, visualizes results, and is integrated with different Bioconductor packages [http://bioconductor.org/packages/PathoStat/]. Pavian is an R package which incorporates Shiny and D3.js components to enable interactive analysis of results for metagenomic classification tools such as Kraken (Breitwieser & Salzberg, 2016). Panviz is a tool for exploring annotated microbial genomics data sets based on D3.js libraries [http://bioconductor.org/packages/PanVizGenerator/]. Krona is a web-based tool for metagenomics visualization that provides a sunburst diagram to navigate the feature space (Ondov et al., 2011). VAMPS is a web-service that provides a JavaScript and PHP based metagenomics visualization toolkit of datasets uploaded by researchers (Huse et al., 2014). Anvi’o is a multiomics platform that supports analysis along with custom JavaScript visualizations (Eren et al. 2015).

### Future Work

An avenue for continued research in this area is visualization of whole metagenome shotgun sequencing data. This will involve both navigation of the feature taxonomy tree as well as exploration of the genes for each bacterial feature. This will be a useful visualization as strain level analysis is expected to be useful for research and clinical applications. Immediate problems in this case are merging with a functional hierarchy for genes in different features and aligning in a track viewer the different genomes present in a sample.

## Acknowledgements

This work was partially supported by NIH grant RO1GM114267 to JW, JK, HCB, NIH grant U54DK102556 to JW, VF, AM and HCB, and a US National Science Foundation Graduate Research Fellowship (award DGE0750616) to JNP.

